# Placental microRNA Expression Associates with Birthweight through Control of Adipokines: Results from Two Independent Cohorts

**DOI:** 10.1101/2020.04.28.067025

**Authors:** Elizabeth M Kennedy, Karen Hermetz, Amber Burt, Todd M Everson, Maya Deyssenroth, Ke Hao, Jia Chen, Margaret R Karagas, Dong Pei, Devin C Koestler, Carmen J Marsit

## Abstract

MicroRNAs are non-coding RNAs that regulate gene expression post-transcriptionally. In the placenta, the master regulator of fetal growth and development, microRNAs shape the basic processes of trophoblast biology and specific microRNA have been associated with fetal growth. To comprehensively assess the role of microRNAs in placental function and fetal development, we have performed small RNA sequencing to profile placental microRNAs from two independent mother-infant cohorts: the Rhode Island Child Health Study (n=225) and the New Hampshire Birth Cohort Study (n=317). We modeled microRNA counts on infant birthweight percentile (BWP) in each cohort, while accounting for race, sex, parity and technical factors, using negative binomial generalized linear models. We identified microRNAs that were differentially expressed (DEmiRs) with BWP at false discovery rate (FDR) less than 0.05 in both cohorts. hsa-miR-532-5p (miR-532) was positively associated with BWP in both cohorts. By integrating parallel whole transcriptome and small RNA sequencing in the RICHS cohort, we identified putative targets of miR-532. These targets are enriched for pathways involved in adipogenesis, adipocytokine signaling, energy metabolism and hypoxia response, and included Leptin, which we further demonstrated to have decreasing expression with increasing BWP, particularly in male infants. Overall, we have shown a robust and reproducible association of miR-532 with BWP, which could influence BWP through regulation of adipocytokines Leptin and Adiponectin.

## INTRODUCTION

MicroRNAs are 21-25 base pair non-coding RNAs that regulate gene expression post-transcriptionally. In humans, microRNAs, as part of the RNA-induced silencing complex (miRISC), target mRNA transcripts based on sequence complementarity to the 3’ untranslated region of the mRNA. The miRISC complex inhibits the translation of target mRNAs through one of three mechanisms: 1) through direct cleavage of the mRNA, 2) recruiting mRNA degradation machinery or 3) blocking translational machinery. MicroRNAs are thought to regulate more than 50% of human genes ^1–4^.

The placenta is the master regulator of early development. During implantation, the trophectoderm, which gives rise to the placenta, invades through the extracellular matrix of the decidualized endometrium into the uterine wall^5–7^. Implantation is followed by placentation, in which the trophectoderm differentiates and proliferates into a network of branching villi – the functional units of the placenta which facilitate the exchange of gases, nutrients and waste for the growing embryo^8,9^. Poor placental function is the most common cause of intrauterine growth restriction, which is associated with perinatal morbidity, mortality and long-term health outcomes. Previous research, primarily conducted in cell lines and model systems, suggests placental microRNAs are pivotal to placental development and function by targeting genes that control trophoblast proliferation and differentiation, apoptosis, invasion, cellular metabolism, as well as vasculo- and angio-genesis (reviewed in ^4^).

A number of studies have investigated the role of microRNA expression in placental function and birth outcomes ^10–29^. Several have studied links between microRNA expression and low birthweight or intrauterine growth restriction (IUGR)^21–29^. Some of these studies have performed targeted analyses of candidate microRNAs, using single^27^ or multiplex qPCR^24^. Many used microarray technology to target a larger number of microRNAs^15,23,25,26^. Most of these studies had sample sizes below 100 ^21,22,24–28^. Although some of these studies divided their own cohorts into discovery and validation subsets^22,23,25,26^, none of them replicated their findings in independently collected cohorts. From these studies, it is clear that microRNA play an essential role in placental biology and function; and that adverse pregnancy outcomes are associated with differential expression of various microRNAs. However, well powered, data-driven and reproducible epidemiologic studies investigating the association of placental microRNAs and birthweight are lacking.

To assess the role of all expressed microRNAs in placental function and fetal health at term, we have utilized small RNA sequencing to profile placental microRNAs from two independent mother-infant cohorts: the New Hampshire Birth Cohort Study (NHBCS; n=317) and Rhode Island Child Health Study (RICHS; n=225). We examined the relationships between microRNAs and birthweight percentiles (BWP) independently in RICHS and NHBCS, and evaluated consistency across these cohorts, then used integrated parallel -omics data to characterize a microRNA that associates with BWP at term, as well as its putative mRNA targets.

## METHODS

### Cohorts

#### The New Hampshire Birth Cohort Study (NHBCS)

was initiated in 2009 and is an ongoing study comprised of a cohort of mother-infant pairs. Pregnant women between 18 and 45 years of age were recruited from the study’s participating prenatal care clinics in New Hampshire, USA. Women were included in the cohort if their primary source of drinking water was from an unregulated residential well, they had resided in the same household since their last menstrual period and had no plans to move before delivery. All participants provided written informed consent in accordance with the requirements of the Committee for the Protection of Human Subjects, the Institutional Review Board (IRB) of Dartmouth College. In this study, NHBCS participants were singleton pregnancies recruited between February 2012 and September 2013. Placental microRNA transcript abundance data were provided for this cohort (*n* = 317). Questionnaires and medical record abstraction were utilized to collect sociodemographic, lifestyle, and anthropometric data (**Table 1**).

**Table 1.** Cohort demographics.

|  | **NHBCS (n = 317)** | **RICHs (n = 225)** |
| --- | --- | --- |
| **MATERNAL CHARACTERISTICS** |  |  |
| **Primigravida - % (n)** | 27% (85) | 24% (54) |
| **INFANT CHARACTERISTICS** |  |  |
| **Race (White) - % (n)** | 100% (317) | 73% (165) |
| **Gestational Age (wks) - mean (range)** | 40 (31 – 42) | 39 (37 – 41) |
| **Birthweight (g) - mean (range)** | 3440 (1380 – 5216) | 3540 (2030 – 5465) |
| **Birthweight Group** |  |  |
| SGA - % (n) | 4% (14) | 16% (36) |
| AGA - % (n) | 86% (273) | 56% (127) |
| LGA - % (n) | 10% (30) | 28% (62) |
| **Sex (Female) - % (n)** | 50% (159) | 49% (111) |

#### The Rhode Island Child Health Study (RICHS)

enrolled mother-infant pairs from the Women & Infants Hospital in Providence, Rhode Island, between September 2010 and February 2013. Mothers were at least 18 years of age, had no life-threatening conditions, and delivered singletons free of congenital/chromosomal abnormalities at or after 37 weeks of gestation. Infants who were born small for gestational age (≤ 10th BWP) or large for gestational age (≥ 90th BWP) were oversampled. Infants who were adequate for gestational age (between the 10th and 90th BWPs), and matched for gestational age and maternal age were coincidentally enrolled. All participants provided written informed consent and all protocols were approved by the IRBs at the Women & Infants Hospital of Rhode Island and Emory University, respectively. Data provided by this study include placental microRNA transcript abundance (n=230) and total RNA transcript abundance (RNA-seq, n=199). Interviewer-administered questionnaires were utilized to collect sociodemographic and lifestyle data. Structured medical record review was used to collect anthropometric and medical history data.

### Data collection

#### Anthropomorphic measures

Both of the studies have parallel demographic and anthropomorphic measures for mothers and newborns, as well as placental microRNA transcript abundance. Birthweight (g), head circumference (cm), and birth length (cm) were abstracted from medical records. *z*-Scores were calculated for each, standardized by gestational age and infant sex, via Fenton growth curves ^30^. Infants with standardized BWP below the 10th percentile are classified as small for gestational age (SGA); those above the 90th percentile as large for gestational age (LGA); and those between the 10th and 90th percentile as adequate for gestational age (AGA). SGA, AGA and LGA are collectively referred to as birthweight groups (BWG). Mother’s BMI was calculated from height and weight collected from medical records.

#### Additional study covariates

Fetal sex was obtained from medical records. Fetal race was self-reported and coded as “white” or “not white”.

#### Tissue collection

The same placental sampling protocol was used by both cohorts. Fetal placental samples were collected within two hours of birth; sections were obtained two centimeters (cm) from the umbilical cord and free of maternal decidua. Collected tissue was immediately placed in RNA later solution (Life Technologies, Grand Island, NY, USA) and stored at 4 °C for at least 72 hours. Subsequently, tissue segments were blotted dry and stored at −80 °C.

#### NHBCS microRNA isolation and sequencing

Total RNA was extracted from fixed placenta using the Qiagen miRNeasy Mini Kit and a TissueLyser LT (Qiagen, Frederick, MD, USA) following manufacturer’s protocol. Briefly, 25-35 mg of placental tissue was placed in a 2 ml round bottom tube with 700 ul of Qiazol Lysing Reagent and one 5 mm stainless steel bead. The tissue was homogenized on the TissueLyser LT for 2 minutes at 40 Hz. The resulting homogenate was processed with the Qiagen miRNeasy Mini Kit and eluted in 30 µl RNase-free water. The RNA was quantitated on a NanoDrop 2000 (Thermo Fisher, Waltham, MA, USA) and quality checked on Agilent Bioanalyzer using the Agilent RNA 6000 Nano kit (Agilent, Santa Clara, CA, USA). Single end, 1 x 75 bp next generation sequencing of placental microRNA was performed by Qiagen Genomic Services (Frederick, Maryland).

#### RICHS microRNA isolation and sequencing

Total RNA was extracted from fixed, pulverized placenta using the Qiagen miRNeasy Mini Kit and a TissueLyser LT (Qiagen, Germantown, MD, USA) following manufacturer’s protocol. Briefly, 25-35 mg of frozen, powdered placental tissue was placed in a 2 ml round bottom tube with 700 ul of Qiazol Lysing Reagent and one 5 mm stainless steel bead. The tissue was homogenized in a pre-chilled tube holder on the TissueLyser LT for two, 5-minute cycles at 30 Hz. The resulting homogenate was processed with the Qiagen miRNeasy Mini Kit with on-column DNAse digestion and eluted in 50 µl RNase-free water. The RNA was quantitated on a NanoDrop (Thermo Fisher, Waltham, MA, USA) and quality checked on Agilent Bioanalyzer using the Agilent RNA 6000 Nano kit (Agilent, Santa Clara, CA, USA). Single end, 1 x 50 bp next generation sequencing of placental microRNA was performed by Omega Bioservices (Norcross, Georgia).

#### RICHS total RNA isolation, sequencing, processing and quality control

Total RNA was extracted from fixed, pulverized placenta using the RNeasy Mini Kit (Qiagen) and stored at −80°C until analysis. We quantified RNA using a Nanodrop Spectrophotometer (Thermo Scientific), assessed RNA integrity via Agilent Bioanalyzer (Agilent), removed ribosomal RNA with a Ribo-Zero Kit ^31^, converted to cDNA using random hexamers (Thermo Scientific), and performed transcriptome-wide 50 bp single-end RNA sequencing via the HiSeq 2500 platform (Illumina) ^32^. Initial quality control was performed on raw reads using the FastQC software. Reads that passed the quality control metrics were mapped to the human reference genome (hg19) using the Spliced Transcripts Alignment to a Reference (STAR) aligner^33^, with common SNPs in the reference genome masked prior to alignment. Raw reads have been deposited at the NCBI sequence read archive (SRP095910).

#### smallRNA-Seq Processing and Quality Control

Raw FASTQ reads obtained from a total of 552 (RICHS 230, NHBCS 322) samples were subject to adaptor trimming with *cutadapt* v1.16^34^. For RICHS reads, the 3’ adaptor sequence were trimmed (TGGAATTCTCGGGTGCCAAGG) and then four bases were trimmed from each end of the read following vendor’s recommendation (BIOO scientific, Austin TX). For NHBCS reads, the 3’ adaptor sequence (AACTGTAGGCACCATCAAT) was trimmed based on vendor’s recommendation (Qiagen). After adaptor trimming, *fastQC* v0.11.5 was used to process the trimmed reads and QC results were aggregated using *MultiQC* v1.5 for visualization^35^. One sample from NHBCS failed QC and was removed. Then we used trimmed reads and *miRDeep2* to quantify microRNA^36^. In short, *miRDeep2* was used to first perform alignment using *bowtie1* with human genome hg38^37^. The ‘Quantifier’ module in *miRDeep2* was used to obtain raw counts of microRNAs with *miRBase* version 22 ^38^.

#### Sample filtering and transcript filtering

Raw counts were imported into *DESeq2* for normalization and differential expression analysis. microRNAs with less than one count per million in more than 10 percent of samples were removed. Five samples were excluded from RICHS microRNA data set because of missing race variable. Four samples in the NHBCS microRNA data set were missing birthweight and were excluded. Of the 2,656 microRNA transcripts that mapped in both RICHS and NHBCS, 802 and 777 remained after filtering, respectively. Of the 50,810 total RNA transcripts that mapped in RICHS, 17,607 remained.

#### Normalization

Filtered microRNA and RNA raw counts were imported to DESeq2 for normalization and differential expression analysis. For all data sets, parametric estimates of dispersion were calculated, and the median ratio method was used to estimate size factors for normalization for modeling with *DESeq2*^39^. Normalized counts were exported from *DESeq2* for surrogate variable analysis. Count matrices normalized by the Variance Stabilizing Transformation (equivalent to normalization and log_2_-transformation) were exported from *DESeq2* for the microRNA and RNA datasets in RICHS, for correlation analysis.

### Statistical analyses

#### Surrogate variable analysis

To adjust for unmeasured confounding effects, such as cell-type heterogeneity and other unmeasured sources of technical variation, we estimated surrogate variables from the normalized transcript reads (for microRNA and total RNA counts) via the *svaseq* function in the *sva* package ^41,42^. In the *svaseq* function, the iteratively re-weighted least squares algorithm was used to estimate surrogate variables based on empirically-derived control transcripts. The full model (mod argument) used for *svaseq* differed by application. The null model (mod0 argument) used all covariates except BWP. For RICHS and NHBCS microRNA differential expression analysis, one surrogate variable was estimated. For RICHS mRNA differential expression analysis, three surrogate variables were estimated.

#### Differential expression analyses

Transcript counts from the microRNA and total RNA sequencing datasets were modeled using a negative binomial generalized linear model with significance testing for differentially expressed transcripts via Wald tests in *DESeq2*^40^. The microRNA transcripts were regressed on infant BWP (or BWG for follow-up analyses), with covariates for race (RICHS analyses only), sex, primigravida; and technical variables. In RICHS microRNA analysis, RIN (RNA integrity number) and sequencing lane number were included in the model as technical variables. In NHBCS microRNA analysis, RIN and sequencing pool were included in the model as technical variables. Surrogate variables were also included as covariates in our regression models (discussed in previous section). We considered microRNAs with a false discovery rate less than 5% to be primary differentially expressed (DEmiRs) with BWP.

In RICHS RNA analysis, RNA transcript abundances were regressed on BWG, with covariates for race, sex, primigravida, sequencing batch and three surrogate variables. Contrasts were used to estimate the log_2_ fold change in the transcript for SGA and LGA, both relative to AGA. To assess the possible interaction of sex and birthweight, RNA transcripts were regressed on the interaction term of infant sex and BWG, with the same covariates listed previously. For these follow-up analyses, candidate transcripts with p-values less than 0.05 for either LGA or SGA were considered significant.

#### Differential expression comparison

Results from the RICHS and NHBCS microRNA differential expression analyses were filtered to the 719 microRNAs common to both analyses. The Pearson correlation of the RICHS and NHBCS differential expression analyses Wald statistics was calculated using the *cor.test* function in R.

#### Identifying mRNA targets of hsa-miR-532-5p

Potential mRNA targets hsa-miR-532-5p (miR-532) were identified from mirDIP, an online database of human microRNA-target predictions^43^. mirDIP integrates microRNA target prediction across 30 different resources, providing nearly 152 million human microRNA-target predictions. Using the individual source ranking and confidence measures, mirDIP assigns a unified rank and confidence score using the quadratic function. For this analysis, the predictions with the top 5% confidence scores were returned^43^. The resulting miR-532-target mRNA pairs were carried forward in the analysis.

#### Putative target filtering

microRNA and total RNA counts were correlated using the Pearson method with the R library *psych*^44^. miR-532-target mRNA pairs that were negatively correlated (FDR < 0.05) were carried forward in the analysis and the associated mRNA were considered putative miR-532 target mRNAs. Finally, putative miR532 target mRNAs that were significantly associated with BWG (P<0.05) in RICHS, were selected for pathway analysis.

#### Pathway analysis

Pathway over-representation analysis was conducted in consensuspathDB^45^, which aggregates data from 12 separate pathway analysis databases. For each pathway gene set, consensus path DB calculates a p-value according to the hypergeometric test for the genes in both the miR-532 putative target genes and the pathway gene set. All mRNAs not filtered for low reads were included as the background, or null distribution, for the test.

## RESULTS

This study analyzed data from 317 mother–infant pairs from the New Hampshire Birth Cohort (NHBCS) and 225 mother–infant pairs from the Rhode Island Child Health Study (RICHS). The demographics of the participants for this study shown in Table 1. A few important differences exist between the two cohorts. By design, RICHS contains a greater proportion of SGA and LGA newborns than NHBCS (**Table 1**). The majority of placentae in both studies were from full term pregnancies and the median gestational ages were similar, NHBCS has a larger range of gestational ages than RICHS. Lastly, the RICHS cohort is more racially diverse than the NHBCS cohort (**Table 1**).

To analyze the associations between BWP and placental microRNA expression, we performed differential expression analysis. Within each cohort and for each microRNA transcript individually, transcript abundances were regressed on corresponding newborn BWP in *DESeq2*^40^, while also adjusting for primigravida, infant sex, race (white/not), technical covariates (RNA integrity number, sequencing lane, library pool) and one surrogate variable^42^. In NHBCS, 777 microRNA transcripts passed filtering, 7 had an FDR <0.05 and 2 were significant with a Bonferroni threshold (p-value < 7e-05, **Figure 1a, Table S1**). Of the 802 microRNA transcripts that passed filtering in RICHS, 11 had an FDR < 0.05 and 3 passed a strict Bonferroni threshold (p-value < 7.2e-05, **Figure 1b, Table S2**).

**Figure 1.**
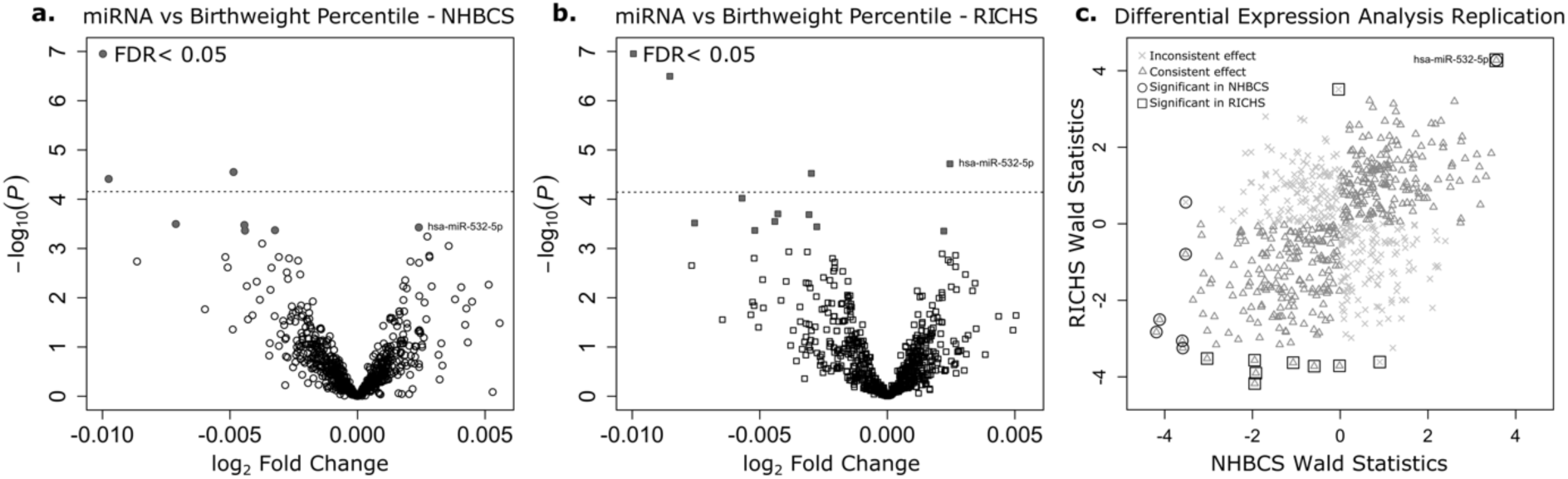
microRNA association with birthweight percentile. Volcano plots illustrate the results of the differential expression analyses for the New Hampshire Birth Cohort Study (round points; **a**) and the Rhode Island Child Health Study (square points; **b**), respectively. On the y and x axes, -log_10_(p-values) in the association of each microRNA with BWP and effect estimates, or the log_2_ fold change in each microRNA per one percent change in BWP, are shown respectively. The horizontal dashed lines represent the Bonferroni threshold (log_10_[0.05/M], where M is the number of microRNAs tested). **a)** 7 microRNAs are significantly (FDR < 0.05) associated with BWP in NHBCS (grey filled shapes), with two microRNAs significant after Bonferroni correction. **b)** 11 microRNAs are significantly (FDR < 0.05) associated with BWP in RICHS (grey filled shapes), with three microRNAs significant after Bonferroni correction. **c)** Wald statistics for the association of each microRNA with BWP from the cohort-level differential expression analyses were plotted and labeled to indicate if they were: consistent in direction of effect (empty triangle), significant in RICHS (square), or significant in NHBCS (circle).

Of the microRNA transcripts that passed filtering in RICHS and NHBCS, 719 were common. Among the primary DEMirs, miR-532 had a significant positive association with BWP in both NHBCS (log_2_ fold change = 2.4e-3; log_2_ standard error: 6.7e-4; FDR = 0.04; **Figure 1a**) and RICHS (log_2_ fold change = 2.4e-3; log_2_ standard error: 5.7e-4; FDR = 6.6e-3; **Figure 1b**). Although there was little overlap between the microRNAs associated with BWP in the two cohorts, the Wald statistics from the cohort-level differential expression analysis were correlated (Pearson’s coefficient = 0.35, P = 1.3e-21), indicating that the NHBCS and RICHS differential expression analyses produced similar results despite sampling differences (**Figure 1c**).

In order to examine whether the placentas of babies born very large or very small exhibited different expression levels of miR-532, we also performed a differential expression analysis where microRNA transcript abundances were regressed on BWG (SGA/AGA/LGA). The positive correlation of miR-532 and birthweight was still evident in the extremes of birthweight groupings (**Figure 2**). When compared to the AGA group, there was a 0.12 and 0.24 log_2_ fold change decrease in miR-532 in the SGA group, for RICHS and NHBCS, respectively. There is an apparent increase of miR-532 from the AGA to LGA groups (0.07 and 0.1 log_2_ fold change, for RICHS and NHBCS), although the lower bounds of the confidence intervals crossed the null (**Figure 2**).

**Figure 2.**
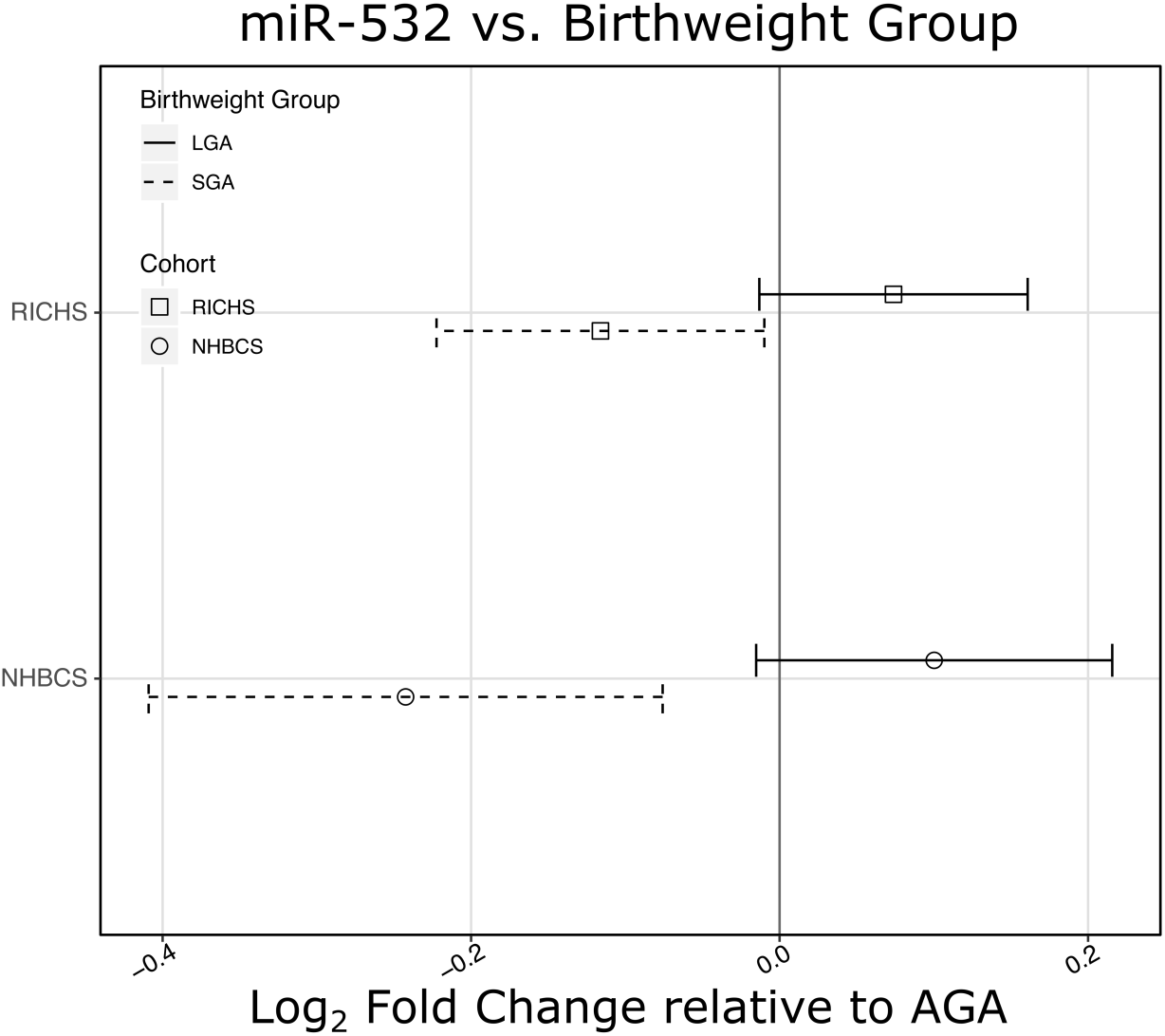
miR-532 associates with birthweight group. Estimates of log_2_ fold change for SGA and LGA, both relative to AGA, illustrate a significant reduction of miR-532 in placentae from SGA newborns and an apparent increase in miR-532 in LGA newborns for NHBCS and RICHS. Birthweight groups, small (SGA) and large (LGA) for gestational age are represented by dashed and solid lines, respectively.

### miR-532 is associated with the expression of impactful targets

Bioinformatic targets of miR-532 were collected from the microRNA Data Integration Portal (mirDIP)^43^. Bioinformatic predictions are known to generate false positives. mirDIP predicted 1,883 targets of miR-532. Recent research has indicated that microRNA-guided transcriptional inhibition will result in transcript degradation, leading to less detectable target transcript^1^. In order to enrich our targets with true microRNA-target pairs, we utilized the parallel measures of total RNA and microRNA in the RICHS cohort. mRNA transcripts were considered putative targets of miR-532 if they were bioinformatically predicted targets of miR-532 and if the mRNA’s abundance was negatively correlated with miR-532 transcript abundance in RICHS (**Table S3**). Using these criteria, we found 56 putative miR-532 targets. We further curated our putative targets of miR-532 by requiring that they be nominally associated (p-value < 0.05) with birthweight in the RICHS cohort. These 18 putative miR-532 target mRNAs used in pathway analysis (**Figure 3, Table S3**).

**Figure 3.**
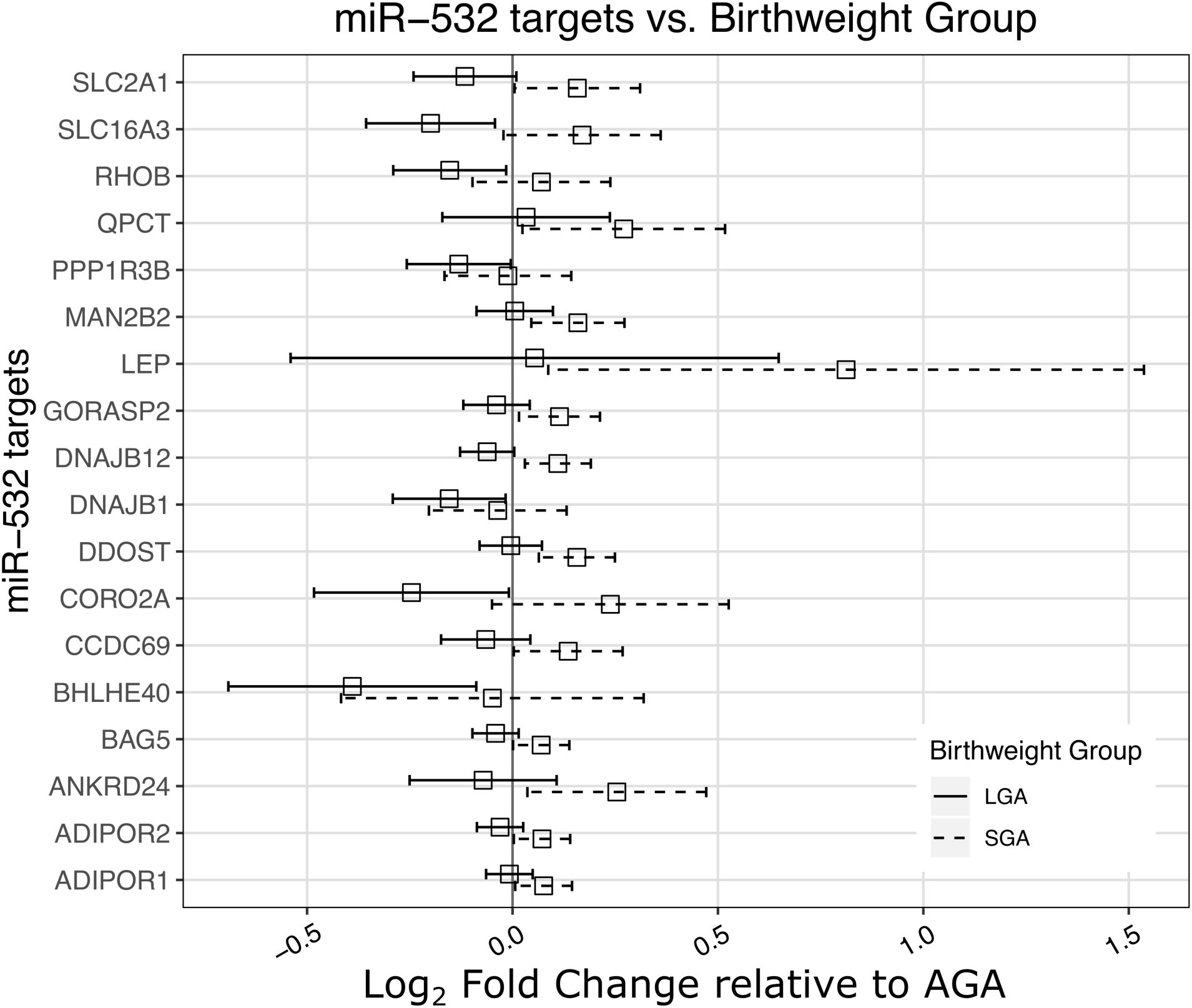
miR-532 target transcript abundance association with LGA or SGA, relative to AGA. Log_2_ fold change of miR-532 putative target transcript abundance in placentae from SGA and LGA newborns, relative to AGA newborns.

The eighteen putative miR-532 target genes were tested for pathway overrepresentation with consensuspathdb, against all of the genes that passed QC in the RICHS whole transcriptome RNA-seq analysis^45^. Pathways related to adipogenesis, adipocytokine signaling and energy metabolism were over-represented among miR-532 putative targets. These enrichment results are primarily driven by putative miR-532 target mRNA transcripts: Leptin (*LEP)*, Adiponectin Receptor 1 (*ADIPOR1)*, Adiponectin Receptor 2 (*ADIPOR2)* and Solute Carrier Family 2 Member 1 (*SLC2A1, GLUT1)* (**Figure 4**).

**Figure 4.**
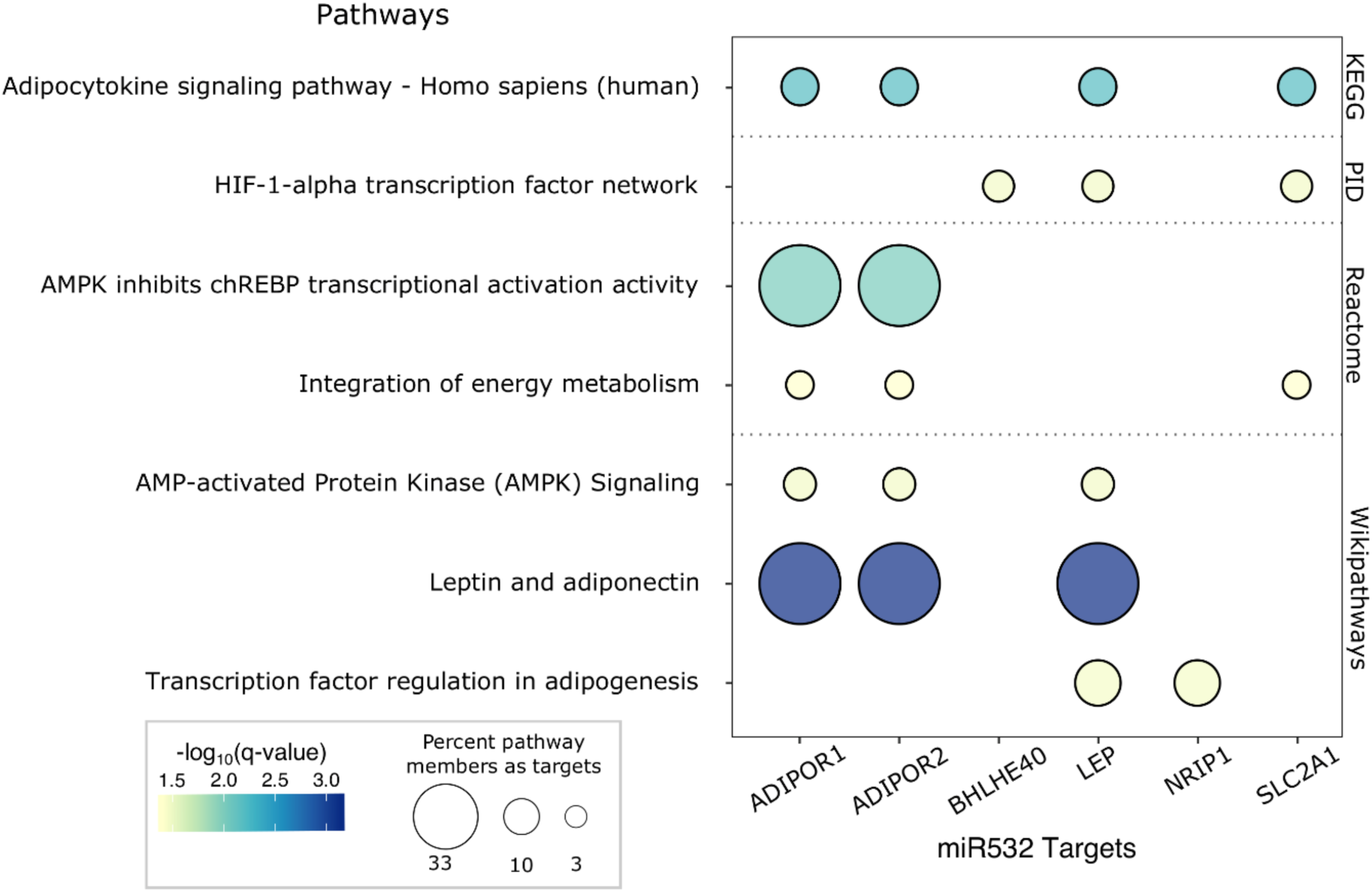
Pathway analysis of miR-532 putative targets. Select consensus path DB results are presented in a bubble plot. Dot color represents -log_10_(FDR). Dot size indicates the percentage of pathway members that are represented by miR-532 targets. Pathway sources are indicated on the right y-axis.

### Sex bias in the associations of LEP with birthweight

We have previously shown a sexually dimorphic association of DNA methylation at the *LEP* promoter with birthweight in RICHS placentae^46^. Five of the 18 putative miR-532 target mRNAs, including *LEP* had significant interactions between sex and BWG (**Table S4**). Because of the enrichment of adipokine pathways in miR-532 target mRNAs and our previous findings regarding *LEP*, we used a stratified analysis to assess sex-specific differences. We found that *LEP* transcript abundance is negatively associated with BWP in males, but not associated with birthweight in females (**Figure 5b**). There is a one-fold decrease in *LEP* from AGA to LGA and a reciprocal (though not significant) one-fold increase in *LEP* from AGA to SGA group. For females in this analysis, there are apparent (though not significant) increases in *LEP* in SGA and LGA groups, relative to the AGA (**Figure 5a**).

**Figure 5.**
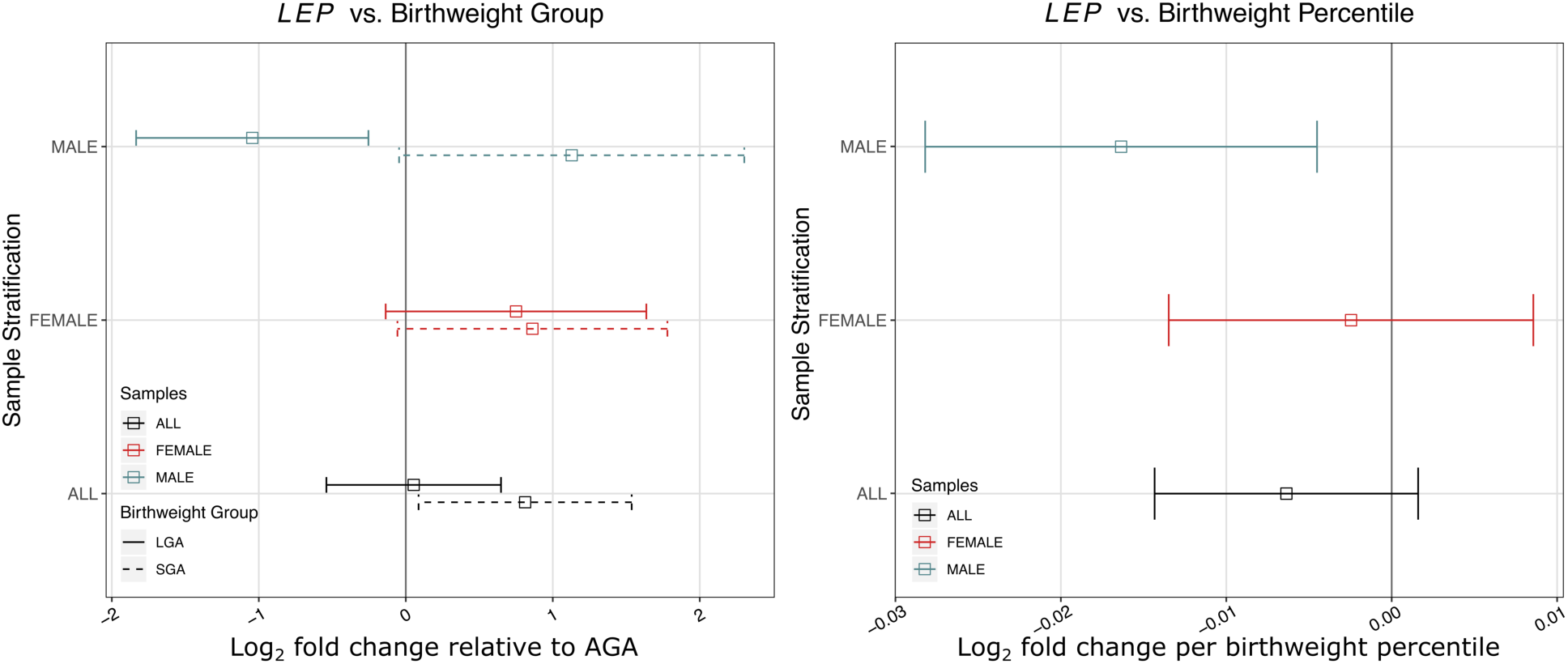
Sex-specific association of birthweight percentile and group with placental leptin transcript abundance in RICHS. **a)** log_2_ fold change of *LEP* abundance in placentae from SGA and LGA newborns, relative to AGA newborns. **b)** Estimates of log_2_ fold change per one BWP. Color represents sample stratification: black (all placentae), red (female placentae) and blue (male placentae).

## DISCUSSION

In this study, we have used an integrated, multi-omics approach to describe the microRNAs that associate with birthweight in human term placenta. We performed these analyses across two mother-infant cohorts, NHBCS and RICHS, to compare consistency in the overall miRNA relationships, which revealed an overall correlation in the relationships and one miRNA that achieved an FDR < 0.05 in both. We found evidence of differentially expressed microRNAs in both cohorts, with one microRNA found to be similarly and statistically significantly differentially expressed in both cohorts. We utilized parallel whole and small RNA-seq in RICHS to narrow microRNA target mRNAs to only those that were negatively correlated with miR-532 transcript abundance. We further filtered miR-532 target mRNAs to only those displaying a nominally significant association with birthweight in the RICHS cohort. These miR-532 mRNA targets were used in a pathway analysis, revealing enrichments of adipogenesis, adipocytokine signaling and energy metabolism pathways.

The roles of microRNAs are dependent on the mRNAs they regulate, and these interactions are tissue and context specific. Few non-targeted microRNA studies in placenta exist; fewer studying microRNA and birthweight. To our knowledge, miR-532 expression in placenta has not been previously associated with birthweight. In two studies of birthweight, miR-532 was chosen as an endogenous control, due to stable transcript levels^11,23^. However, in the study from Timofeeva, et al, miR-532 was differentially expressed in the placenta of women with preeclampsia, suggesting it may not be a suitable stable control ^11^. In this study, miR-532 was differentially expressed with birth weight percentiles and when comparing SGA and LGA to AGA in two independent cohorts, with similar estimates of effect in both cohorts, indicating that this association is robust and reproducible.

While we focus on miR-532, which had the strongest evidence for an association with birth weight, several other microRNAs yielded strong associations with BWP in either NHBCS or RICHS, some of which were consistent with the findings of other studies (**Table S5**). Among microRNA with an FDR < 0.1 in NHBCS, three were also found in similar studies (hsa-miR-193b-3p^21,28^, hsa-miR-181a^23^, hsa-miR-101-5p^21^). In RICHS, nine microRNAs were consistent with previous findings (hsa-miR-520a-3p^21^, hsa-miR-193b-5p^21^, hsa-miR-210-3p^21^, hsa-miR-193b-3p^21,28^, hsa-miR-335-3p^28^, hsa-miR-515-5p^21,22^, hsa-miR-519d-5p^21,22,26^, hsa-miR-365b-3p^21^, hsa-miR-365a-3p^21^). It is reassuring that despite different sampling schemes, study designs and nucleic acid technologies - we see an overlap in these findings even though they did not meet the statistical threshold of FDR < 0.05 in both cohorts for our study.

Bioinformatic gene target prediction and integration with mRNA target transcript abundances indicate that gene targets of miR-532 in our samples were enriched in adipogenesis and energy metabolism pathways. The adipokines leptin and adiponectin modulate insulin sensitivity, metabolism, and energy homeostasis in adulthood^47,48^. While their roles in placenta are less clear, both of these hormones seem to be involved in placental development and function^49,50^. *LEP* has a placenta-specific enhancer and in placenta, leptin has autocrine functions related to angiogenesis, hypoxia response and endocrine functions including mobilization of maternal fat^51^. Unlike leptin, adiponectin is not produced by the placenta ^52^. However through the placental expression of adiponectin receptors, maternal adiponectin might limit fetal growth by inhibiting the production of placental lactogen, chorion gonadotropin and progesterone in trophoblast cells^52,53^. Both of these hormones have been linked to gestational diseases like preeclampsia^54–58^, gestational diabetes^58–61^ and fetal outcomes like intrauterine growth restriction^46,58,59,62–66^. While earlier work did find positive associations between placental leptin expression and birthweight^59,60,67^, several studies since have found a negative correlation – which is in line with our findings^63,65,68–72^.

In adulthood, leptin is known to be responsive to gonadal steroids, resulting in a sexually dimorphic response to leptin signaling^73^. We have previously reported sexually dimorphic methylation patterns in the *LEP* promoter^46^. Lesseur *et al* found that RICHS male placentae had higher levels of promoter methylation than female placentae^46^. Assuming the canonical negative association of promoter DNA methylation and gene expression, those results are consistent with our own, in which the mean *LEP* transcript counts for male placentae were lower than for female placentae. To our knowledge, we are the first research study to find a dimorphic association of *LEP* expression with birthweight in placenta. Cagnacci et. al report that leptin in amniotic fluid at 16 weeks is inversely correlated with fetal growth in females and not correlated with any growth parameter in males^72^. This study makes a case for sexual dimorphic negative association between leptin and fetal growth, but in the opposite sex as in our study. We found that *LEP* expression inversely correlated to BWP in newborn males, and was not associated with BWP in females. The findings described in the Cagnacci study are at a much earlier time in gestation and in a different tissue. At 16 weeks of gestation, the placental production of leptin is low^74^ and the amniotic fluid is supplied primarily by maternal serum^75^, suggesting that leptin concentration could be a representation of the mother’s leptin status and unrelated to placental leptin^72^.

To our knowledge, this is the largest study yet to examine the relationships between human placental microRNAs and birthweight and the only study in which results are replicated in an independent cohort. Our results suggest a role for miR-532 in the regulation of fetal growth through control of genes that control energy metabolism. However, our findings should be interpreted within the context of this study’s limitations.

This is an observational study in which RNA was assayed from term placentae. Thus, we cannot conclude that our results are representative of microRNA associations throughout development, and we cannot determine the directionality of these observed relationships since BWP and microRNA were measured cross-sectionally. We adjusted for likely confounders in our study, but cannot rule out the possibility that unmeasured or residual confounding remains in our analysis. Two potential confounders are maternal BMI and gestational weight gain. These variables were not included in the analysis, because they were not available for the NHBCS cohort at the time of this analysis. Sensitivity analysis suggests that the addition of maternal BMI and gestational weight gain to our models would have little to no effect on our conclusions, though some of our top findings are sensitive to one or both (**Tables S6-S11, Figures S1 and S2**). To limit statistical confounding due to unknown confounding (e.g. differing cellular composition between individuals or population stratification), our models are adjusted using surrogate variable analysis. This is a data-driven approach that may incorrectly estimate confounding elements, adding variability to our model. Recent research indicates that SVA is one of the more robust and reliable methods in studies such as ours^76^. Lastly, both cohorts utilized in this study consisted predominantly of healthy white mothers from the New England region of the United States, potentially limiting the generalizability of our findings.

## CONCLUSION

MicroRNAs are an important regulatory mechanism in gene expression. The activity of microRNAs has been linked to the development and function of placentae throughout gestation, making them essential to fetal development. Here we have shown that the transcript abundance of miR-532 is associated with birthweight in independent cohorts. Putative targets of miR-532 are involved in angiogenesis, energy metabolism and endocrine signaling. Although targets of miR-532 are already established as regulators of placental development and function, ours is the first study to report the association of miR-532 and birthweight.

## SUPPLEMENTAL FIGURES

S1 – NHBCS Sensitivity Analysis

S2 – RICHS Sensitivity Analysis

## SUPPLEMENTAL TABLES

S1 – NHBCS Differential Expression Analysis results

S2 – RICHS Differential Expression Analysis results

S3 – Putative miR-532 target association with BWG and empirical correlation with miR-532

S4 – Putative miR-532 target vs BWG:sex interaction

S5 – Results comparison to previous studies

S6 – NHBCS Gestational weight gain category Sensitivity Analysis

S7 – NHBCS pre-pregnancy BMI Sensitivity Analysis

S8 – NHBCS gestational weight gain and pre-pregnancy BMI sensitivity analysis

S9 – RICHS Gestational weight gain category Sensitivity Analysis

S10 – RICHS pre-pregnancy BMI Sensitivity Analysis

S11 – RICHS gestational weight gain and pre-pregnancy BMI sensitivity analysis

## References

1. Bartel DP. Metazoan MicroRNAs. Cell. 2018 Mar 22;173(1):20–51.

2. Huberdeau MQ, Simard MJ. A guide to microRNA-mediated gene silencing. FEBS J. 2019;286(4):642–652.

3. Pasquinelli AE. MicroRNAs and their targets: recognition, regulation and an emerging reciprocal relationship. Nat Rev Genet. 2012 Apr;13(4):271–282.

4. Hayder H, O’Brien J, Nadeem U, Peng C. MicroRNAs: crucial regulators of placental development. Reproduction. 2018 Jun 1;155(6):R259–R271.

5. Brosens I, Robertson WB, Dixon HG. The physiological response of the vessels of the placental bed to normal pregnancy. J Pathol Bacteriol. 1967 Apr;93(2):569–579. PMID: 6054057

6. Meekins JW, Pijnenborg R, Hanssens M, McFadyen IR, van Asshe A. A study of placental bed spiral arteries and trophoblast invasion in normal and severe pre-eclamptic pregnancies. Br J Obstet Gynaecol. 1994 Aug;101(8):669–674. PMID: 7947500

7. Regnault TRH, Galan HL, Parker TA, Anthony RV. Placental development in normal and compromised pregnancies--a review. Placenta. 2002 Apr;23 Suppl A:S119–129. PMID: 11978069

8. Red-Horse K, Zhou Y, Genbacev O, Prakobphol A, Foulk R, McMaster M, Fisher SJ. Trophoblast differentiation during embryo implantation and formation of the maternal-fetal interface. J Clin Invest. American Society for Clinical Investigation; 2004 Sep 15;114(6):744–754. PMID: 15372095

9. Kaufmann P, Mayhew TM, Charnock-Jones DS. Aspects of Human Fetoplacental Vasculogenesis and Angiogenesis. II. Changes During Normal Pregnancy. Placenta. 2004 Feb 1;25(2):114–126.

10. Guo D, Jiang H, Chen Y, Yang J, Fu Z, Li J, Han X, Wu X, Xia Y, Wang X, Chen L, Tang Q, Wu W. Elevated microRNA-141-3p in placenta of non-diabetic macrosomia regulate trophoblast proliferation. EBioMedicine. 2018 Dec 1;38:154–161.

11. Timofeeva AV, Gusar VA, Kan NE, Prozorovskaya KN, Karapetyan AO, Bayev OR, Chagovets VV, Kliver SF, Iakovishina DYu, Frankevich VE, Sukhikh GT. Identification of potential early biomarkers of preeclampsia. Placenta. 2018 Jan 1;61:61–71.

12. Wu L, Song W, Xie Y, Hu L, Hou X, Wang R, Gao Y, Zhang J, Zhang L, Li W, Zhu C, Gao Z, Sun Y. miR-181a-5p suppresses invasion and migration of HTR-8/SVneo cells by directly targeting IGF2BP2. Cell Death Dis. 2018 Jan 16;9(2):1–14.

13. Enquobahrie DA, Abetew DF, Sorensen TK, Willoughby D, Chidambaram K, Williams MA. Placental microRNA expression in pregnancies complicated by preeclampsia. Am J Obstet Gynecol. 2011 Feb 1;204(2):178.e12–178.e21.

14. Vashukova ES, Glotov AS, Fedotov PV, Efimova OA, Pakin VS, Mozgovaya EV, Pendina AA, Tikhonov AV, Koltsova AS, Baranov VS. Placental microRNA expression in pregnancies complicated by superimposed pre-eclampsia on chronic hypertension. Mol Med Rep. Spandidos Publications; 2016 Jul 1;14(1):22–32.

15. Yang S, Li H, Ge Q, Guo L, Chen F. Deregulated microRNA species in the plasma and placenta of patients with preeclampsia. Mol Med Rep. Spandidos Publications; 2015 Jul 1;12(1):527–534.

16. Niu Z, Han T, Sun X, Luan L, Gou W, Zhu X. MicroRNA-30a-3p is overexpressed in the placentas of patients with preeclampsia and affects trophoblast invasion and apoptosis by its effects on IGF-1. Am J Obstet Gynecol. 2018 Feb 1;218(2):249.e1–249.e12.

17. Jiang F, Li J, Wu G, Miao Z, Lu L, Ren G, Wang X. Upregulation of microRNA-335 and microRNA-584 contributes to the pathogenesis of severe preeclampsia through downregulation of endothelial nitric oxide synthase. Mol Med Rep. Spandidos Publications; 2015 Oct 1;12(4):5383–5390.

18. Zhu X, Han T, Sargent IL, Yin G, Yao Y. Differential expression profile of microRNAs in human placentas from preeclamptic pregnancies vs normal pregnancies. Am J Obstet Gynecol. 2009 Jun 1;200(6):661.e1–661.e7.

19. Carreras-Badosa G, Bonmatí A, Ortega F-J, Mercader J-M, Guindo-Martínez M, Torrents D, Prats-Puig A, Martinez-Calcerrada J-M, de Zegher F, Ibáñez L, Fernandez-Real J-M, Lopez-Bermejo A, Bassols J. Dysregulation of Placental miRNA in Maternal Obesity Is Associated With Pre- and Postnatal Growth. J Clin Endocrinol Metab. 2017 01;102(7):2584–2594. PMID: 28368446

20. Li J, Song L, Zhou L, Wu J, Sheng C, Chen H, Liu Y, Gao S, Huang W. A MicroRNA Signature in Gestational Diabetes Mellitus Associated with Risk of Macrosomia. Cell Physiol Biochem. 2015;37(1):243–252. PMID: 26302821

21. Awamleh Z, Gloor GB, Han VKM. Placental microRNAs in pregnancies with early onset intrauterine growth restriction and preeclampsia: potential impact on gene expression and pathophysiology. BMC Med Genomics. 2019 Jun 27;12(1):91.

22. Higashijima A, Miura K, Mishima H, Kinoshita A, Jo O, Abe S, Hasegawa Y, Miura S, Yamasaki K, Yoshida A, Yoshiura K, Masuzaki H. Characterization of placenta-specific microRNAs in fetal growth restriction pregnancy. Prenat Diagn. 2013;33(3):214–222.

23. Rahman ML, Liang L, Valeri L, Su L, Zhu Z, Gao S, Mostofa G, Qamruzzaman Q, Hauser R, Baccarelli A, Christiani DC. Regulation of birthweight by placenta-derived miRNAs: evidence from an arsenic-exposed birth cohort in Bangladesh. Epigenetics. 2018;13(6):573–590. PMCID: PMC6140906

24. Wang D, Na Q, Song W-W, Song G-Y. Altered Expression of miR-518b and miR-519a in the Placenta is Associated with Low Fetal Birth Weight. Am J Perinatol. 2014 Oct;31(9):729–734.

25. Meng M, Cheng YKY, Wu L, Chaemsaithong P, Leung MBW, Chim SSC, Sahota DS, Li W, Poon LCY, Wang CC, Leung TY. Whole genome miRNA profiling revealed miR-199a as potential placental pathogenesis of selective fetal growth restriction in monochorionic twin pregnancies. Placenta. 2020 Mar 1;92:44–53.

26. Thamotharan S, Chu A, Kempf K, Janzen C, Grogan T, Elashoff DA, Devaskar SU. Differential microRNA expression in human placentas of term intra-uterine growth restriction that regulates target genes mediating angiogenesis and amino acid transport. PLOS ONE. 2017 May 2;12(5):e0176493.

27. Huang L, Shen Z, Xu Q, Huang X, Chen Q, Li D. Increased levels of microRNA-424 are associated with the pathogenesis of fetal growth restriction. Placenta. 2013 Jul 1;34(7):624–627.

28. Östling H, Kruse R, Helenius G, Lodefalk M. Placental expression of microRNAs in infants born small for gestational age. Placenta. 2019 Jun 1;81:46–53.

29. Guo L, Tsai SQ, Hardison NE, James AH, Motsinger-Reif AA, Thames B, Stone EA, Deng C, Piedrahita JA. Differentially expressed microRNAs and affected biological pathways revealed by modulated modularity clustering (MMC) analysis of human preeclamptic and IUGR placentas. Placenta. 2013 Jul 1;34(7):599–605.

30. Fenton TR, Kim JH. A systematic review and meta-analysis to revise the Fenton growth chart for preterm infants. BMC Pediatr. 2013 Apr 20;13:59. PMCID: PMC3637477

31. Huang R, Jaritz M, Guenzl P, Vlatkovic I, Sommer A, Tamir IM, Marks H, Klampfl T, Kralovics R, Stunnenberg HG, Barlow DP, Pauler FM. An RNA-Seq strategy to detect the complete coding and non-coding transcriptome including full-length imprinted macro ncRNAs. PloS One. 2011;6(11):e27288. PMCID: PMC3213133

32. Bentley DR, Balasubramanian S, Swerdlow HP, Smith GP, Milton J, Brown CG, Hall KP, Evers DJ, Barnes CL, Bignell HR, Boutell JM, Bryant J, Carter RJ, Keira Cheetham R, Cox AJ, Ellis DJ, Flatbush MR, Gormley NA, Humphray SJ, Irving LJ, Karbelashvili MS, Kirk SM, Li H, Liu X, Maisinger KS, Murray LJ, Obradovic B, Ost T, Parkinson ML, Pratt MR, Rasolonjatovo IMJ, Reed MT, Rigatti R, Rodighiero C, Ross MT, Sabot A, Sankar SV, Scally A, Schroth GP, Smith ME, Smith VP, Spiridou A, Torrance PE, Tzonev SS, Vermaas EH, Walter K, Wu X, Zhang L, Alam MD, Anastasi C, Aniebo IC, Bailey DMD, Bancarz IR, Banerjee S, Barbour SG, Baybayan PA, Benoit VA, Benson KF, Bevis C, Black PJ, Boodhun A, Brennan JS, Bridgham JA, Brown RC, Brown AA, Buermann DH, Bundu AA, Burrows JC, Carter NP, Castillo N, Chiara E Catenazzi M, Chang S, Neil Cooley R, Crake NR, Dada OO, Diakoumakos KD, Dominguez-Fernandez B, Earnshaw DJ, Egbujor UC, Elmore DW, Etchin SS, Ewan MR, Fedurco M, Fraser LJ, Fuentes Fajardo KV, Scott Furey W, George D, Gietzen KJ, Goddard CP, Golda GS, Granieri PA, Green DE, Gustafson DL, Hansen NF, Harnish K, Haudenschild CD, Heyer NI, Hims MM, Ho JT, Horgan AM, Hoschler K, Hurwitz S, Ivanov DV, Johnson MQ, James T, Huw Jones TA, Kang G-D, Kerelska TH, Kersey AD, Khrebtukova I, Kindwall AP, Kingsbury Z, Kokko-Gonzales PI, Kumar A, Laurent MA, Lawley CT, Lee SE, Lee X, Liao AK, Loch JA, Lok M, Luo S, Mammen RM, Martin JW, McCauley PG, McNitt P, Mehta P, Moon KW, Mullens JW, Newington T, Ning Z, Ling Ng B, Novo SM, O’Neill MJ, Osborne MA, Osnowski A, Ostadan O, Paraschos LL, Pickering L, Pike AC, Pike AC, Chris Pinkard D, Pliskin DP, Podhasky J, Quijano VJ, Raczy C, Rae VH, Rawlings SR, Chiva Rodriguez A, Roe PM, Rogers J, Rogert Bacigalupo MC, Romanov N, Romieu A, Roth RK, Rourke NJ, Ruediger ST, Rusman E, Sanches-Kuiper RM, Schenker MR, Seoane JM, Shaw RJ, Shiver MK, Short SW, Sizto NL, Sluis JP, Smith MA, Ernest Sohna Sohna J, Spence EJ, Stevens K, Sutton N, Szajkowski L, Tregidgo CL, Turcatti G, Vandevondele S, Verhovsky Y, Virk SM, Wakelin S, Walcott GC, Wang J, Worsley GJ, Yan J, Yau L, Zuerlein M, Rogers J, Mullikin JC, Hurles ME, McCooke NJ, West JS, Oaks FL, Lundberg PL, Klenerman D, Durbin R, Smith AJ. Accurate whole human genome sequencing using reversible terminator chemistry. Nature. 2008 Nov 6;456(7218):53–59. PMCID: PMC2581791

33. Dobin A, Davis CA, Schlesinger F, Drenkow J, Zaleski C, Jha S, Batut P, Chaisson M, Gingeras TR. STAR: ultrafast universal RNA-seq aligner. Bioinformatics. 2013 Jan;29(1):15–21. PMCID: PMC3530905

34. Martin M. Cutadapt removes adapter sequences from high-throughput sequencing reads. EMBnet.journal. 2011 May 2;17(1):10–12.

35. Ewels P, Magnusson M, Lundin S, Käller M. MultiQC: summarize analysis results for multiple tools and samples in a single report. Bioinforma Oxf Engl. 2016 01;32(19):3047–3048. PMCID: PMC5039924

36. Friedländer MR, Mackowiak SD, Li N, Chen W, Rajewsky N. miRDeep2 accurately identifies known and hundreds of novel microRNA genes in seven animal clades. Nucleic Acids Res. 2012 Jan;40(1):37–52. PMCID: PMC3245920

37. Langmead B, Trapnell C, Pop M, Salzberg SL. Ultrafast and memory-efficient alignment of short DNA sequences to the human genome. Genome Biol. 2009;10(3):R25. PMCID: PMC2690996

38. Griffiths-Jones S, Saini HK, van Dongen S, Enright AJ. miRBase: tools for microRNA genomics. Nucleic Acids Res. 2008 Jan;36(Database issue):D154–158. PMCID: PMC2238936

39. Anders S, Huber W. Differential expression analysis for sequence count data. Genome Biol. 2010 Oct 27;11(10):R106.

40. Leek JT, Johnson WE, Parker HS, Jaffe AE, Storey JD. The sva package for removing batch effects and other unwanted variation in high-throughput experiments. Bioinforma Oxf Engl. 2012 Mar 15;28(6):882–883. PMCID: PMC3307112

41. Leek JT. svaseq: removing batch effects and other unwanted noise from sequencing data. Nucleic Acids Res. 2014 Dec 1;42(21):e161–e161.

42. Love MI, Huber W, Anders S. Moderated estimation of fold change and dispersion for RNA-seq data with DESeq2. Genome Biol. 2014 Dec 5;15(12):550.

43. Tokar T, Pastrello C, Rossos AEM, Abovsky M, Hauschild A-C, Tsay M, Lu R, Jurisica I. mirDIP 4.1—integrative database of human microRNA target predictions. Nucleic Acids Res. 2018 Jan 4;46(Database issue):D360–D370. PMCID: PMC5753284

44. Revelle WR. psych: Procedures for Personality and Psychological Research. 2017 [cited 2020 Mar 26]; Available from: https://www.scholars.northwestern.edu/en/publications/psych-procedures-for-personality-and-psychological-research

45. Kamburov A, Pentchev K, Galicka H, Wierling C, Lehrach H, Herwig R. ConsensusPathDB: toward a more complete picture of cell biology. Nucleic Acids Res. 2011 Jan;39(Database issue):D712–717. PMCID: PMC3013724

46. Lesseur C, Armstrong DA, Paquette AG, Koestler DC, Padbury JF, Marsit CJ. Tissue-specific Leptin promoter DNA methylation is associated with maternal and infant perinatal factors. Mol Cell Endocrinol. 2013 Dec 5;381(1–2):160–167. PMCID: PMC3795868

47. Seeley RJ, Woods SC. Monitoring of stored and available fuel by the CNS: implications for obesity. Nat Rev Neurosci. Nature Publishing Group; 2003 Nov;4(11):901–909.

48. Stefan N, Stumvoll M. Adiponectin--its role in metabolism and beyond. Horm Metab Res Horm Stoffwechselforschung Horm Metab. 2002 Sep;34(9):469–474. PMID: 12384822

49. Schanton M, Maymó JL, Pérez-Pérez A, Sánchez-Margalet V, Varone CL. Involvement of leptin in the molecular physiology of the placenta. Reproduction. 2018 Jan 1;155(1):R1–R12.

50. Costa MA. The endocrine function of human placenta: an overview. Reprod Biomed Online. 2016 Jan 1;32(1):14–43.

51. Bi S, Gavrilova O, Gong D-W, Mason MM, Reitman M. Identification of a Placental Enhancer for the Human Leptin Gene. J Biol Chem. American Society for Biochemistry and Molecular Biology; 1997 Nov 28;272(48):30583–30588. PMID: 9374555

52. McDonald EA, Wolfe MW. Adiponectin attenuation of endocrine function within human term trophoblast cells. Endocrinology. 2009 Sep;150(9):4358–4365. PMID: 19520781

53. Caminos JE, Nogueiras R, Gallego R, Bravo S, Tovar S, García-Caballero T, Casanueva FF, Diéguez C. Expression and Regulation of Adiponectin and Receptor in Human and Rat Placenta. J Clin Endocrinol Metab. Oxford Academic; 2005 Jul 1;90(7):4276–4286.

54. Weiwei T, Haiyan Y, Juan C, Xiaodong W, Weibo C, Rong Z. Expressions of Adiponectin Receptors in Placenta and Their Correlation With Preeclampsia. Reprod Sci. 2009 Jul 1;16(7):676–684.

55. Meller M, Qiu C, Kuske BT, Abetew DF, Muy-Rivera M, Williams MA. Adipocytokine expression in placentas from pre-eclamptic and chronic hypertensive patients. Gynecol Endocrinol. Taylor & Francis; 2006 Jan 1;22(5):267–273.

56. Hytinantti T, Koistinen HA, Koivisto VA, Karonen S-L, Rutanen E-M, Andersson S. Increased leptin concentration in preterm infants of pre-eclamptic mothers. Arch Dis Child - Fetal Neonatal Ed. 2000 Jul 1;83(1):F13–F16. PMID: 10873164

57. Laivuori H, Gallaher MJ, Collura L, Crombleholme WR, Markovic N, Rajakumar A, Hubel CA, Roberts JM, Powers RW. Relationships between maternal plasma leptin, placental leptin mRNA and protein in normal pregnancy, pre-eclampsia and intrauterine growth restriction without pre-eclampsia. Mol Hum Reprod. 2006 Sep 1;12(9):551–556.

58. Pérez-Pérez A, Toro A, Vilariño-García T, Maymó J, Guadix P, Dueñas JL, Fernández-Sánchez M, Varone C, Sánchez-Margalet V. Leptin action in normal and pathological pregnancies. J Cell Mol Med. 2018;22(2):716–727.

59. Lea RG, Howe D, Hannah LT, Bonneau O, Hunter L, Hoggard N. Placental leptin in normal, diabetic and fetal growth-retarded pregnancies. Mol Hum Reprod. 2000 Aug 1;6(8):763–769.

60. Lepercq J, Cauzac M, Lahlou N, Timsit J, Girard J, Auwerx J, Mouzon SH. Overexpression of placental leptin in diabetic pregnancy: a critical role for insulin. Diabetes. 1998 May 1;47(5):847–850. PMID: 9588462

61. Uzelac PS, Li X, Lin J, Neese LD, Lin L, Nakajima ST, Bohler H, Lei Z. Dysregulation of Leptin and Testosterone Production and Their Receptor Expression in the Human Placenta with Gestational Diabetes Mellitus. Placenta. 2010 Jul 1;31(7):581–588.

62. Wang J, Shang L-X, Dong X, Wang X, Wu N, Wang S-H, Zhang F, Xu L-M, Xiao Y. Relationship of adiponectin and resistin levels in umbilical serum, maternal serum and placenta with neonatal birth weight. Aust N Z J Obstet Gynaecol. 2010;50(5):432–438.

63. Lazo-de-la-Vega-Monroy M-L, González-Domínguez MI, Zaina S, Sabanero M, Daza-Benítez L, Malacara JM, Barbosa-Sabanero G. Leptin and its Receptors in Human Placenta of Small, Adequate, and Large for Gestational Age Newborns. Horm Metab Res. 2017 May;49(5):350–358.

64. Denisova EI, Kozhevnikova VV, Bazhan NM, Makarova EN. Sex-specific effects of leptin administration to pregnant mice on the placentae and the metabolic phenotypes of offspring. FEBS Open Bio. 2020 Jan;10(1):96–106. PMCID: PMC6943234

65. Schrey S, Kingdom J, Baczyk D, Fitzgerald B, Keating S, Ryan G, Drewlo S. Leptin is differentially expressed and epigenetically regulated across monochorionic twin placenta with discordant fetal growth. Mol Hum Reprod. 2013 Nov 1;19(11):764–772.

66. Tzschoppe AA, Struwe E, Dörr HG, Goecke TW, Beckmann MW, Schild RL, Dötsch J. Differences in gene expression dependent on sampling site in placental tissue of fetuses with intrauterine growth restriction. Placenta. 2010 Mar 1;31(3):178–185.

67. Hoggard N, Haggarty P, Thomas L, Lea RG. Leptin expression in placental and fetal tissues: does leptin have a functional role? Biochem Soc Trans. 2001 May 1;29(2):57–63.

68. McCarthy C, Cotter FE, McElwaine S, Twomey A, Mooney EE, Ryan F, Vaughan J. Altered gene expression patterns in intrauterine growth restriction: Potential role of hypoxia. Am J Obstet Gynecol. 2007 Jan 1;196(1):70.e1–70.e6.

69. McMinn J, Wei M, Schupf N, Cusmai J, Johnson EB, Smith AC, Weksberg R, Thaker HM, Tycko B. Unbalanced Placental Expression of Imprinted Genes in Human Intrauterine Growth Restriction. Placenta. 2006 Jun 1;27(6):540–549.

70. Struwe E, Berzl G, Schild R, Blessing H, Drexel L, Hauck B, Tzschoppe A, Weidinger M, Sachs M, Scheler C, Schleussner E, Dötsch J. Microarray analysis of placental tissue in intrauterine growth restriction. Clin Endocrinol (Oxf). 2010;72(2):241–247.

71. Sabri A, Lai D, D’Silva A, Seeho S, Kaur J, Ng C, Hyett J. Differential Placental Gene Expression in Term Pregnancies Affected by Fetal Growth Restriction and Macrosomia. Fetal Diagn Ther. Karger Publishers; 2014;36(2):173–180. PMID: 24685769

72. Cagnacci A, Arangino S, Caretto S, Mazza V, Volpe A. Sexual dimorphism in the levels of amniotic fluid leptin in pregnancies at 16 weeks of gestation: Relation to fetal growth. Eur J Obstet Gynecol Reprod Biol. 2006 Jan 1;124(1):53–57.

73. Power ML, Schulkin J. Sex differences in fat storage, fat metabolism, and the health risks from obesity: possible evolutionary origins. Br J Nutr. Cambridge University Press; 2008 May;99(5):931–940.

74. Forhead AJ, Fowden AL. The hungry fetus? Role of leptin as a nutritional signal before birth. J Physiol. 2009 Mar 15;587(Pt 6):1145–1152. PMCID: PMC2674987

75. Underwood MA, Gilbert WM, Sherman MP. Amniotic Fluid: Not Just Fetal Urine Anymore. J Perinatol. Nature Publishing Group; 2005 May;25(5):341–348.

76. van Rooij J, Mandaviya PR, Claringbould A, Felix JF, van Dongen J, Jansen R, Franke L, ‘t Hoen PAC, Heijmans B, van Meurs JBJ, BIOS consortium. Evaluation of commonly used analysis strategies for epigenome- and transcriptome-wide association studies through replication of large-scale population studies. Genome Biol. 2019 Nov 14;20(1):235.

