## Supplemental Tables and Figures for "Placental microRNA Expression Associates with Birthweight through Control of Adipokines: Results from Two Independent Cohorts"

### SUPPLEMENTAL FIGURES

NHBCS Sensitivity analysis: top 100 DEmiRs from DEA

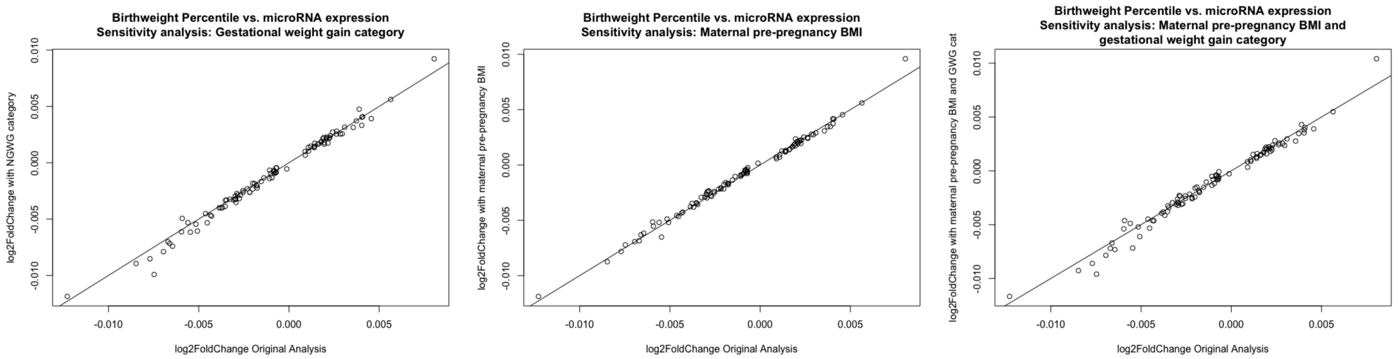


Figure S1. Sensitivity Analyses. Effects for the top 100 results (by p-value) from the original analysis are plotted against the effects from the a) gestational weight gain, b) pre-pregnancy BMI and c) combined gestational weight gain and BMI sensitivity analyses of the same 100 microRNAs.

RICHS Sensitivity Analysis: top 100 DEmiRs from DEA

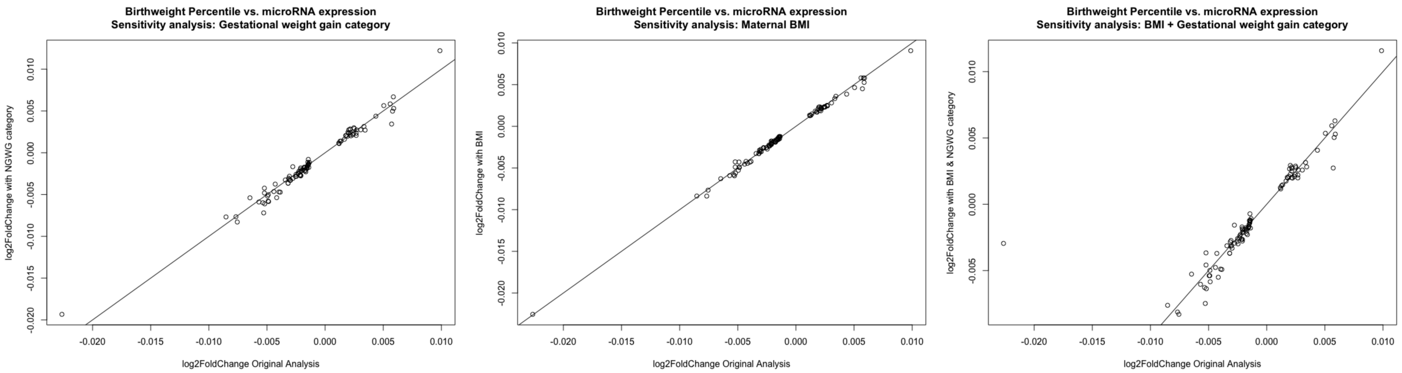


Figure S2. Sensitivity Analyses. Effects for the top 100 results (by p-value) from the original analysis are plotted against the effects from the a) gestational weight gain, b) pre-pregnancy BMI and c) combined gestational weight gain and BMI sensitivity analyses of the same 100 microRNAs.

**SUPPLEMENTAL TABLES**

Table S1. NHBCS Differential Expression Analysis results (shorter version)

| | baseMean | log2FoldChange | lfcSE | stat | pvalue | padj |
|---|---|---|---|---|---|---|
| hsa-miR-629-3p | 37.706 | -0.005 | 0.001 | -4.189 | 0.000 | 0.014 |
| hsa-miR-155-5p | 1110.267 | -0.010 | 0.002 | -4.115 | 0.000 | 0.014 |
| hsa-miR-1246 | 65.321 | -0.007 | 0.002 | -3.600 | 0.000 | 0.044 |
| hsa-miR-146b-5p | 3397.645 | -0.004 | 0.001 | -3.588 | 0.000 | 0.044 |
| hsa-miR-532-5p | 6177.514 | 0.002 | 0.001 | 3.558 | 0.000 | 0.044 |
| hsa-miR-624-5p | 194.505 | -0.003 | 0.001 | -3.523 | 0.000 | 0.044 |
| hsa-miR-146a-5p | 5756.189 | -0.004 | 0.001 | -3.519 | 0.000 | 0.044 |
| hsa-miR-503-5p | 6627.305 | 0.003 | 0.001 | 3.445 | 0.001 | 0.051 |
| hsa-miR-29b-1-5p | 81.333 | -0.004 | 0.001 | -3.354 | 0.001 | 0.063 |
| hsa-miR-125b-2-3p | 115.993 | 0.004 | 0.001 | 3.324 | 0.001 | 0.064 |
| hsa-let-7c-5p | 6088.727 | 0.003 | 0.001 | 3.195 | 0.001 | 0.073 |
| hsa-miR-449a | 59.828 | -0.003 | 0.001 | -3.177 | 0.001 | 0.073 |
| hsa-miR-146b-3p | 24.878 | -0.005 | 0.002 | -3.176 | 0.001 | 0.073 |
| hsa-miR-542-3p | 24438.726 | 0.003 | 0.001 | 3.173 | 0.002 | 0.073 |
| hsa-miR-627-5p | 109.707 | -0.003 | 0.001 | -3.158 | 0.002 | 0.073 |
| hsa-miR-576-3p | 227.129 | -0.002 | 0.001 | -3.127 | 0.002 | 0.073 |
| hsa-let-7e-5p | 35878.302 | 0.002 | 0.001 | 3.116 | 0.002 | 0.073 |
| hsa-miR-375-3p | 219.936 | -0.009 | 0.003 | -3.115 | 0.002 | 0.073 |
| hsa-miR-1287-5p | 156.019 | 0.002 | 0.001 | 3.098 | 0.002 | 0.073 |
| hsa-miR-193b-3p | 1927.447 | -0.005 | 0.002 | -3.033 | 0.002 | 0.084 |
| hsa-miR-181a-2-3p | 2535.049 | -0.003 | 0.001 | -3.029 | 0.002 | 0.084 |
| hsa-miR-101-5p | 297.650 | -0.003 | 0.001 | -2.963 | 0.003 | 0.099 |
| hsa-miR-22-5p | 2586.019 | -0.002 | 0.001 | -2.932 | 0.003 | 0.105 |
| hsa-miR-1278 | 33.782 | -0.003 | 0.001 | -2.866 | 0.004 | 0.124 |
| hsa-miR-542-5p | 2625.080 | 0.002 | 0.001 | 2.847 | 0.004 | 0.126 |
| hsa-miR-181b-2-3p | 58.342 | -0.004 | 0.001 | -2.826 | 0.005 | 0.130 |
| hsa-miR-6507-5p | 16.909 | 0.005 | 0.002 | 2.779 | 0.005 | 0.141 |
| hsa-miR-4521 | 42.075 | -0.004 | 0.002 | -2.759 | 0.006 | 0.141 |
| hsa-miR-450a-1-3p | 652.451 | 0.002 | 0.001 | 2.756 | 0.006 | 0.141 |
| hsa-miR-99a-5p | 4901.873 | 0.003 | 0.001 | 2.754 | 0.006 | 0.141 |
| hsa-miR-301b-3p | 193.381 | 0.004 | 0.001 | 2.741 | 0.006 | 0.141 |
| hsa-miR-450b-5p | 4164.003 | 0.002 | 0.001 | 2.724 | 0.006 | 0.144 |
| hsa-miR-154-3p | 359.644 | -0.003 | 0.001 | -2.702 | 0.007 | 0.150 |
| hsa-miR-335-3p | 8822.412 | 0.002 | 0.001 | 2.604 | 0.009 | 0.194 |
| hsa-miR-873-5p | 66.141 | 0.004 | 0.001 | 2.550 | 0.011 | 0.215 |
| hsa-miR-548aq-3p | 178.773 | -0.004 | 0.002 | -2.543 | 0.011 | 0.215 |
| hsa-miR-660-5p | 11905.131 | 0.002 | 0.001 | 2.539 | 0.011 | 0.215 |
| hsa-miR-548e-3p | 98.926 | -0.002 | 0.001 | -2.516 | 0.012 | 0.220 |
| hsa-miR-135b-5p | 14972.865 | 0.004 | 0.002 | 2.512 | 0.012 | 0.220 |
| hsa-miR-582-3p | 83.498 | 0.003 | 0.001 | 2.498 | 0.013 | 0.224 |
| hsa-miR-642a-5p | 6.263 | -0.009 | 0.003 | -2.482 | 0.013 | NA |
| hsa-miR-29b-3p | 108050.005 | -0.002 | 0.001 | -2.449 | 0.014 | 0.250 |

Table S2. RICHS Differential Expression Analysis results (shorter version)

| | baseMean | log2FoldChange | lfcSE | stat | pvalue | padj |
|---|---|---|---|---|---|---|
| hsa-miR-365a-5p | 11.918 | -0.009 | 0.002 | -5.112 | 0.000 | 0.000 |
| hsa-miR-532-5p | 1636.574 | 0.002 | 0.001 | 4.275 | 0.000 | 0.007 |
| hsa-miR-520a-3p | 4490.233 | -0.003 | 0.001 | -4.175 | 0.000 | 0.007 |
| hsa-miR-193b-5p | 50.236 | -0.006 | 0.001 | -3.903 | 0.000 | 0.016 |
| hsa-miR-27a-5p | 239.876 | -0.004 | 0.001 | -3.721 | 0.000 | 0.024 |
| hsa-miR-200b-3p | 244.776 | -0.003 | 0.001 | -3.712 | 0.000 | 0.024 |
| hsa-miR-210-3p | 126.681 | -0.004 | 0.001 | -3.630 | 0.000 | 0.026 |
| hsa-miR-95-3p | 87.869 | -0.008 | 0.002 | -3.612 | 0.000 | 0.026 |
| hsa-miR-29a-3p | 24682.277 | -0.003 | 0.001 | -3.566 | 0.000 | 0.028 |
| hsa-miR-193b-3p | 4351.293 | -0.005 | 0.001 | -3.521 | 0.000 | 0.028 |
| hsa-miR-501-3p | 113.523 | 0.002 | 0.001 | 3.513 | 0.000 | 0.028 |
| hsa-miR-146b-5p | 2177.389 | -0.004 | 0.001 | -3.247 | 0.001 | 0.063 |
| hsa-miR-132-5p | 40.360 | -0.003 | 0.001 | -3.245 | 0.001 | 0.063 |
| hsa-miR-214-5p | 131.280 | 0.002 | 0.001 | 3.220 | 0.001 | 0.064 |
| hsa-miR-335-3p | 4488.993 | 0.003 | 0.001 | 3.200 | 0.001 | 0.064 |
| hsa-miR-181b-2-3p | 9.035 | -0.005 | 0.002 | -3.160 | 0.002 | 0.065 |
| hsa-miR-515-5p | 14822.297 | -0.002 | 0.001 | -3.158 | 0.002 | 0.065 |
| hsa-miR-501-5p | 60.699 | 0.002 | 0.001 | 3.136 | 0.002 | 0.066 |
| hsa-miR-519d-5p | 651.793 | -0.002 | 0.001 | -3.107 | 0.002 | 0.066 |
| hsa-miR-654-5p | 66.854 | 0.002 | 0.001 | 3.105 | 0.002 | 0.066 |
| hsa-miR-1246 | 11.326 | -0.008 | 0.003 | -3.059 | 0.002 | 0.073 |
| hsa-miR-3615 | 39.727 | 0.003 | 0.001 | 3.042 | 0.002 | 0.074 |
| hsa-miR-576-5p | 118.087 | -0.002 | 0.001 | -3.023 | 0.003 | 0.076 |
| hsa-miR-2682-5p | 3.164 | 0.010 | 0.003 | 2.984 | 0.003 | NA |
| hsa-miR-30b-3p | 82.230 | -0.002 | 0.001 | -2.981 | 0.003 | 0.081 |
| hsa-miR-498-3p | 275.293 | -0.002 | 0.001 | -2.977 | 0.003 | 0.081 |
| hsa-miR-363-3p | 698.941 | 0.003 | 0.001 | 2.932 | 0.003 | 0.090 |
| hsa-miR-548x-3p | 5.326 | -0.023 | 0.008 | -2.903 | 0.004 | 0.095 |
| hsa-miR-132-3p | 136.890 | -0.003 | 0.001 | -2.888 | 0.004 | 0.096 |
| hsa-miR-548aq-3p | 9.062 | -0.005 | 0.002 | -2.856 | 0.004 | 0.099 |
| hsa-miR-423-5p | 768.342 | -0.002 | 0.001 | -2.846 | 0.004 | 0.099 |
| hsa-miR-520g-3p | 10727.275 | -0.002 | 0.001 | -2.832 | 0.005 | 0.099 |
| hsa-miR-629-3p | 10.789 | -0.004 | 0.001 | -2.830 | 0.005 | 0.099 |
| hsa-miR-365b-3p | 2542.795 | -0.003 | 0.001 | -2.809 | 0.005 | 0.099 |
| hsa-miR-365a-3p | 2542.692 | -0.003 | 0.001 | -2.809 | 0.005 | 0.099 |
| hsa-miR-511-3p | 11.932 | 0.003 | 0.001 | 2.803 | 0.005 | 0.099 |
| hsa-miR-181a-2-3p | 517.836 | -0.003 | 0.001 | -2.799 | 0.005 | 0.099 |
| hsa-miR-519e-5p | 546.532 | -0.002 | 0.001 | -2.769 | 0.006 | 0.105 |
| hsa-miR-511-5p | 53.018 | 0.003 | 0.001 | 2.748 | 0.006 | 0.109 |
| hsa-miR-31-5p | 633.724 | -0.003 | 0.001 | -2.731 | 0.006 | 0.112 |
| hsa-miR-342-5p | 19.311 | 0.003 | 0.001 | 2.705 | 0.007 | 0.118 |
| hsa-miR-139-3p | 11.766 | 0.003 | 0.001 | 2.686 | 0.007 | 0.120 |

Table S3. Putative miR-532 target association with BWG and empirical correlation with miR-532

| miR-532 Target | Assoc with LGA (P-value) | Assoc with SGA (P-value) | Empirical Correlation with miR-532 |
|---|---|---|---|
| ADIPOR2 | 0.291 | 0.04 | -0.24 |
| MAN2B2 | 0.91 | 0.006 | -0.22 |
| ANKRD24 | 0.433 | 0.022 | -0.22 |
| CORO2A | 0.042 | 0.105 | -0.31 |
| GORASP2 | 0.345 | 0.023 | -0.27 |
| QPCT | 0.751 | 0.031 | -0.26 |
| SLC2A1 | 0.07 | 0.043 | -0.31 |
| DNAJB1 | 0.028 | 0.674 | -0.23 |
| BHLHE40 | 0.011 | 0.794 | -0.32 |
| SLC16A3 | 0.013 | 0.083 | -0.31 |
| RHOB | 0.029 | 0.411 | -0.29 |
| DNAJB12 | 0.067 | 0.007 | -0.22 |
| ADIPOR1 | 0.792 | 0.032 | -0.22 |
| BAG5 | 0.15 | 0.046 | -0.24 |
| PPP1R3B | 0.043 | 0.886 | -0.35 |
| LEP | 0.859 | 0.028 | -0.34 |
| CCDC69 | 0.239 | 0.045 | -0.22 |
| DDOST | 0.913 | 0.001 | -0.24 |

Table S4: Putative miR-532 target vs BWG:sex interaction

| Gene Symbol | P-value | |
|---|---|---|
| | Sex:LGA | Sex:SGA |
| *CORO2A* | 3.05E-04 | 1.17E-01 |
| *AUP1* | 2.29E-02 | 2.41E-01 |
| *SLC2A1* | 3.37E-02 | 9.92E-01 |
| *BAG5* | 8.54E-03 | 7.91E-01 |
| *LEP* | 1.65E-03 | 3.07E-01 |

Table S5: Results comparison to previous studies

NHBCS

| | padj | | |
|---|---|---|---|
| hsa-miR-629-3p | 0.014 | | |
| hsa-miR-155-5p | 0.014 | | |
| hsa-miR-1246 | 0.044 | | |
| hsa-miR-146b-5p | 0.044 | | |
| hsa-miR-532-5p | 0.044 | | |
| hsa-miR-624-5p | 0.044 | | |
| hsa-miR-146a-5p | 0.044 | | |
| hsa-miR-503-5p | 0.051 | | |
| hsa-miR-29b-1-5p | 0.063 | | |
| hsa-miR-125b-2-3p | 0.064 | | |
| hsa-let-7c-5p | 0.073 | | |
| hsa-miR-449a | 0.073 | | |
| hsa-miR-146b-3p | 0.073 | | |
| hsa-miR-542-3p | 0.073 | | |
| hsa-miR-627-5p | 0.073 | | |
| hsa-miR-576-3p | 0.073 | | |
| hsa-let-7e-5p | 0.073 | | |
| hsa-miR-375-3p | 0.073 | | |
| hsa-miR-1287-5p | 0.073 | | |
| hsa-miR-193b-3p | 0.084 | Awamleh | Ostling |
| hsa-miR-181a-2-3p | 0.084 | Rahman | |
| hsa-miR-101-5p | 0.099 | Awamleh | |

RICHS

| | padj | |
|---|---|---|
| hsa-miR-365a-5p | 0.000 | |
| hsa-miR-532-5p | 0.007 | |
| hsa-miR-520a-3p | 0.007 | Awamleh |
| hsa-miR-193b-5p | 0.016 | Awamleh |
| hsa-miR-27a-5p | 0.024 | |
| hsa-miR-200b-3p | 0.024 | |
| hsa-miR-210-3p | 0.026 | Awamleh |

| | | | | |
|---|---|---|---|---|
| hsa-miR-95-3p | 0.026 | | | |
| hsa-miR-29a-3p | 0.028 | | | |
| hsa-miR-193b-3p | 0.028 | Awamleh | Ostling | |
| hsa-miR-501-3p | 0.028 | | | |
| hsa-miR-146b-5p | 0.063 | | | |
| hsa-miR-132-5p | 0.063 | | | |
| hsa-miR-214-5p | 0.064 | | | |
| hsa-miR-335-3p | 0.064 | Ostling | | |
| hsa-miR-181b-2-3p | 0.065 | | | |
| hsa-miR-515-5p | 0.065 | Awamleh | Higashijima | |
| hsa-miR-501-5p | 0.066 | | | |
| hsa-miR-519d-5p | 0.066 | Awamleh | Thamotharan | Higashijima |
| hsa-miR-654-5p | 0.066 | | | |
| hsa-miR-1246 | 0.073 | | | |
| hsa-miR-3615 | 0.074 | | | |
| hsa-miR-576-5p | 0.076 | | | |
| hsa-miR-2682-5p | NA | | | |
| hsa-miR-30b-3p | 0.081 | | | |
| hsa-miR-498-3p | 0.081 | | | |
| hsa-miR-363-3p | 0.090 | | | |
| hsa-miR-548x-3p | 0.095 | | | |
| hsa-miR-132-3p | 0.096 | | | |
| hsa-miR-548aq-3p | 0.099 | | | |
| hsa-miR-423-5p | 0.099 | | | |
| hsa-miR-520g-3p | 0.099 | | | |
| hsa-miR-629-3p | 0.099 | | | |
| hsa-miR-365b-3p | 0.099 | Awamleh | | |
| hsa-miR-365a-3p | 0.099 | Awamleh | | |
| hsa-miR-511-3p | 0.099 | | | |
| hsa-miR-181a-2-3p | 0.099 | | | |

Table S6: NHBCS Gestational weight gain category Sensitivity Analysis (Shorter, with percent change from original model estimates)

| | baseMean | log2FoldChange | lfcSE | stat | pvalue | padj | percent.change.estimate |
|---|---|---|---|---|---|---|---|
| hsa-miR-629-3p | 36.742 | -0.002 | 0.002 | -1.317 | 0.188 | 0.735 | 10.917 |
| hsa-miR-155-5p | 1068.487 | -0.012 | 0.003 | -3.513 | 0.000 | 0.347 | -3.441 |
| hsa-miR-1246 | 69.755 | -0.007 | 0.003 | -2.285 | 0.022 | 0.507 | 8.473 |
| hsa-miR-146b-5p | 3213.496 | -0.005 | 0.002 | -3.025 | 0.002 | 0.398 | 5.760 |
| hsa-miR-532-5p | 5776.427 | 0.002 | 0.001 | 2.186 | 0.029 | 0.535 | 6.560 |
| hsa-miR-624-5p | 183.505 | -0.003 | 0.001 | -2.362 | 0.018 | 0.507 | -0.987 |
| hsa-miR-146a-5p | 5462.815 | -0.005 | 0.002 | -2.297 | 0.022 | 0.507 | -2.329 |
| hsa-miR-503-5p | 6593.084 | 0.004 | 0.001 | 3.018 | 0.003 | 0.398 | -0.523 |
| hsa-miR-29b-1-5p | 83.726 | -0.003 | 0.002 | -1.742 | 0.081 | 0.617 | 6.888 |
| hsa-miR-125b-2-3p | 108.328 | 0.003 | 0.002 | 1.668 | 0.095 | 0.640 | -3.082 |

Table S7: NHBCS pre-pregnancy BMI Sensitivity Analysis (Shorter, with percent change from original model estimates)

| | baseMean | log2FoldChange | lfcSE | stat | pvalue | padj | percent.change.estimate |
|---|---|---|---|---|---|---|---|
| hsa-miR-629-3p | 36.742 | -0.002 | 0.002 | -1.116 | 0.265 | 0.851 | -6.453 |
| hsa-miR-155-5p | 1068.487 | -0.012 | 0.003 | -3.526 | 0.000 | 0.329 | -3.285 |
| hsa-miR-1246 | 69.755 | -0.006 | 0.003 | -2.034 | 0.042 | 0.589 | -4.502 |
| hsa-miR-146b-5p | 3213.496 | -0.005 | 0.002 | -2.764 | 0.006 | 0.558 | -5.271 |
| hsa-miR-532-5p | 5776.427 | 0.002 | 0.001 | 1.899 | 0.058 | 0.644 | -8.725 |
| hsa-miR-624-5p | 183.505 | -0.003 | 0.001 | -2.169 | 0.030 | 0.589 | -11.527 |
| hsa-miR-146a-5p | 5462.815 | -0.005 | 0.002 | -2.358 | 0.018 | 0.589 | -1.267 |
| hsa-miR-503-5p | 6593.084 | 0.003 | 0.001 | 2.865 | 0.004 | 0.558 | -7.474 |
| hsa-miR-29b-1-5p | 83.726 | -0.003 | 0.002 | -1.734 | 0.083 | 0.698 | 5.422 |
| hsa-miR-125b-2-3p | 108.328 | 0.002 | 0.001 | 1.651 | 0.099 | 0.702 | -5.868 |

Table S8: NHBCS gestational weight gain and pre-pregnancy BMI sensitivity analysis (Shorter, with percent change from original model estimates)

| | baseMean | log2FoldChange | lfcSE | stat | pvalue | padj | percent.change.estimate |
|---|---|---|---|---|---|---|---|
| hsa-miR-629-3p | 36.742 | -0.002 | 0.002 | -1.194 | 0.232 | 0.816 | 2.178 |
| hsa-miR-155-5p | 1068.487 | -0.012 | 0.003 | -3.422 | 0.001 | 0.250 | -5.012 |
| hsa-miR-1246 | 69.755 | -0.007 | 0.003 | -2.116 | 0.034 | 0.577 | 1.335 |
| hsa-miR-146b-5p | 3213.496 | -0.005 | 0.002 | -2.869 | 0.004 | 0.524 | 1.040 |
| hsa-miR-532-5p | 5776.427 | 0.002 | 0.001 | 1.971 | 0.049 | 0.595 | -2.600 |
| hsa-miR-624-5p | 183.505 | -0.003 | 0.001 | -2.100 | 0.036 | 0.577 | -11.812 |
| hsa-miR-146a-5p | 5462.815 | -0.004 | 0.002 | -2.253 | 0.024 | 0.572 | -2.988 |
| hsa-miR-503-5p | 6593.084 | 0.003 | 0.001 | 2.789 | 0.005 | 0.524 | -7.200 |
| hsa-miR-29b-1-5p | 83.726 | -0.003 | 0.002 | -1.822 | 0.068 | 0.622 | 13.442 |
| hsa-miR-125b-2-3p | 108.328 | 0.002 | 0.002 | 1.561 | 0.119 | 0.671 | -8.062 |

Table S9: RICHS Gestational weight gain category Sensitivity Analysis (Shorter, with percent change from original model estimates)

| | baseMean | log2FoldChange | lfcSE | stat | pvalue | padj | percent.change.estimate |
|---|---|---|---|---|---|---|---|
| hsa-miR-365a-5p | 11.918 | -0.008 | 0.002 | -4.245 | 0.000 | 0.007 | -9.914 |
| hsa-miR-532-5p | 1636.574 | 0.002 | 0.001 | 3.725 | 0.000 | 0.018 | -5.878 |
| hsa-miR-520a-3p | 4490.233 | -0.003 | 0.001 | -4.307 | 0.000 | 0.007 | 10.766 |
| hsa-miR-193b-5p | 50.236 | -0.006 | 0.002 | -3.721 | 0.000 | 0.018 | 3.472 |
| hsa-miR-27a-5p | 239.876 | -0.004 | 0.001 | -3.014 | 0.003 | 0.070 | -12.528 |
| hsa-miR-200b-3p | 244.776 | -0.003 | 0.001 | -3.123 | 0.002 | 0.066 | -8.501 |
| hsa-miR-210-3p | 126.681 | -0.005 | 0.001 | -3.568 | 0.000 | 0.023 | 5.299 |
| hsa-miR-95-3p | 87.869 | -0.008 | 0.002 | -3.659 | 0.000 | 0.018 | 9.694 |
| hsa-miR-29a-3p | 24682.277 | -0.003 | 0.001 | -3.655 | 0.000 | 0.018 | 10.638 |
| hsa-miR-193b-3p | 4351.293 | -0.006 | 0.002 | -3.854 | 0.000 | 0.018 | 17.709 |

Table S10: RICHS pre-pregnancy BMI Sensitivity Analysis (Shorter, with percent change from original model estimates)

| | baseMean | log2FoldChange | lfcSE | stat | pvalue | padj | percent.change.estimate |
|---|---|---|---|---|---|---|---|
| hsa-miR-365a-5p | 11.918 | -0.008 | 0.002 | -4.938 | 0.000 | 0.001 | -1.870 |
| hsa-miR-532-5p | 1636.574 | 0.002 | 0.001 | 4.053 | 0.000 | 0.012 | -3.688 |
| hsa-miR-520a-3p | 4490.233 | -0.003 | 0.001 | -4.194 | 0.000 | 0.009 | 2.150 |
| hsa-miR-193b-5p | 50.236 | -0.006 | 0.001 | -3.983 | 0.000 | 0.012 | 3.850 |
| hsa-miR-27a-5p | 239.876 | -0.004 | 0.001 | -3.586 | 0.000 | 0.029 | -1.933 |
| hsa-miR-200b-3p | 244.776 | -0.003 | 0.001 | -3.516 | 0.000 | 0.032 | -3.542 |
| hsa-miR-210-3p | 126.681 | -0.005 | 0.001 | -3.685 | 0.000 | 0.026 | 2.907 |
| hsa-miR-95-3p | 87.869 | -0.008 | 0.002 | -3.596 | 0.000 | 0.029 | 1.371 |
| hsa-miR-29a-3p | 24682.277 | -0.003 | 0.001 | -3.317 | 0.001 | 0.049 | -5.656 |
| hsa-miR-193b-3p | 4351.293 | -0.006 | 0.001 | -3.776 | 0.000 | 0.022 | 8.123 |

Table S11: RICHS gestational weight gain and pre-pregnancy BMI sensitivity analysis (Shorter, with percent change from original model estimates)

| | baseMean | log2FoldChange | lfcSE | stat | pvalue | padj | percent.change.estimate |
|---|---|---|---|---|---|---|---|
| hsa-miR-365a-5p | 11.918 | -0.008 | 0.002 | -4.180 | 0.000 | 0.009 | -10.601 |
| hsa-miR-532-5p | 1636.574 | 0.002 | 0.001 | 3.609 | 0.000 | 0.020 | -8.148 |
| hsa-miR-520a-3p | 4490.233 | -0.003 | 0.001 | -4.293 | 0.000 | 0.009 | 11.333 |
| hsa-miR-193b-5p | 50.236 | -0.006 | 0.002 | -3.787 | 0.000 | 0.018 | 6.047 |
| hsa-miR-27a-5p | 239.876 | -0.004 | 0.001 | -2.952 | 0.003 | 0.075 | -13.607 |
| hsa-miR-200b-3p | 244.776 | -0.003 | 0.001 | -3.014 | 0.003 | 0.075 | -11.065 |
| hsa-miR-210-3p | 126.681 | -0.005 | 0.001 | -3.648 | 0.000 | 0.020 | 8.077 |
| hsa-miR-95-3p | 87.869 | -0.008 | 0.002 | -3.633 | 0.000 | 0.020 | 9.874 |
| hsa-miR-29a-3p | 24682.277 | -0.003 | 0.001 | -3.489 | 0.000 | 0.027 | 6.061 |
| hsa-miR-193b-3p | 4351.293 | -0.006 | 0.002 | -4.012 | 0.000 | 0.012 | 22.628 |
